# The most probable ancestral sequence reconstruction yields proteins without systematic bias in thermal stability or activity

**DOI:** 10.1101/2023.02.22.529562

**Authors:** Michael A. Sennett, Brian C. Beckett, Douglas L. Theobald

**Affiliations:** Brandeis University, Department of Biochemistry, Waltham, MA 02453, USA

## Abstract

Ancestral sequence resurrection (ASR) is the inference of extinct biological sequences from extant sequences, the most popular of which are based on probabilistic models of evolution. ASR is becoming a popular method for studying the evolution of enzyme characteristics. The properties of ancestral enzymes are biochemically and biophysically characterized to gain some knowledge regarding the origin of some enzyme property. Current methodology relies on resurrection of the single most probable (SMP) sequence and is systematically biased. Previous theoretical work suggests this will result in a thermostability bias in resurrected SMP sequences, and even the activity, calling into question inferences derived from ancestral protein properties. We experimentally test the potential stability bias hypothesis by resurrecting 40 malate and lactate dehydrogenases. Despite the methodological bias in resurrecting an SMP protein, the measured biophysical and biochemical properties of the SMP protein are not biased in comparison to other, less probable, resurrections. In addition, the SMP protein property seems to be representative of the ancestral probability distribution. Therefore, the conclusions and inferences drawn from the SMP protein are likely not a source of bias.

**Significance:** Ancestral sequence resurrection (ASR) is a powerful tool for: determining how new protein functions evolve; inferring the properties of an environment in which species existed; and protein engineering applications. We demonstrate, using lactate and malate dehydrogenases (L/MDHs), that resurrecting the single most probable sequence (SMP) from a maximum likelihood phylogeny does not result in biased activity and stability relative to sequences sampled from the posterior probability distribution. Previous studies using experimentally measured phenotypes of SMP sequences to make inferences about the environmental conditions and the path of evolution are likely not biased in their conclusions. Serendipitously, we discover ASR is also a valid tool for protein engineering because sampled reconstructions are both highly active and stable.

## Introduction

Ancestral sequence reconstruction (ASR) is the inference of extinct sequences from a set of homologous extant sequences.^1^ Ancestral protein resurrection (APR) is a specific application of ASR typically used to generate proteins in the lab for experimentally testing hypotheses of a protein’s specific physicochemical properties.^2^ APR has been successful as a biochemical tool for several purposes including, demonstrating the thermophilic origins of life, explaining the origin and retention of oligomeric interfaces, revealing key residues for discriminating drug-target binding, and to isolate key residues in the emergence of novel enzyme function.^3–7^ Current state-of-the-art APR methods to reconstruct and resurrect ancestral enzymes use probabilistic models of evolution.^8, 9^ Consequently, there is an inherent uncertainty in the resurrected ancestral sequences that is expected to carry over to conclusions drawn from APR methods.

The most popular programs for ASR use model-based probabilistic inference, by way of maximum likelihood, to generate a phylogeny and predict a distribution of states for an ancestor at each internal node of the phylogeny.^10, 11^ The number of possible sequences inferred is incredibly high because a distribution of nucleotides, codons, or amino acids are predicted at each site in a sequence. Given resource and time constraints, all sequence combinations from the probability distribution for a single node cannot be experimentally characterized to test hypotheses regarding protein evolution. A practical and reasonable solution is to select the single most probable (SMP) state at each site and resurrect that sequence.^12–14^

Resurrecting the SMP ancestor is a reasonable sequence choice to experimentally study because it is the sequence that is expected to have the fewest number of errors from the ‘true’ ancestral sequence.^15–17^ This, of course, is conditional on ancestral state probabilities accurately reflecting the actual uncertainty of the residue and the internal node actually existing. We have demonstrated that the number of correct residues is linearly proportional to the average sequence probability.^18^ Regardless, if selecting the SMP state does indeed result in the fewest sequence errors then we should also expect that it’s physicochemical properties when resurrected is most like the true ancestral protein. However, the concern remains that systematically selecting the most probable residue at each site will impart a property bias in the SMP sequence relative to the true sequence.^19–21^

Selecting a SMP residue ignores the uncertainty in a resurrection and places a disproportional weight toward the SMP residue. As a result, the frequency of inferred ancestral residues for the SMP ancestors has been shown to not reflect their frequency of occurrence observed at the tips.^20^ Protein properties, like activity and stability, are determined by the protein sequence. If the amino acid frequencies in the SMP ancestral sequences are biased, then the SMP protein function is expected to be biased. Further support from computational studies have demonstrated that using the SMP sequence results in biased amino acid composition and stability, and even suggest an activity bias relative to the true ancestral properties.^19^ Their calculations also predict this bias should be eliminated by randomly sampling from the ancestral probability distribution. In light of these theoretical calculations, some experimental observations on resurrected ancestral sequences have been interpreted as evidence in support of biased biochemical SMP sequence properties. For example, a study on the stability of the last common ancestor of mammalian paraoxonases were unexpectedly ~30 °C more stable than their extant counterparts, despite the fact that the internal temperature of an ancestral mammal should be similar to extant mammals.^21^ To mitigate the possibility of bias in their SMP proteins effecting their results experimentalists will typically assemble a preponderance of evidence to demonstrate that their conclusions are robust to sequence uncertainty. For example, some studies will resurrect SMP sequences from alternative phylogenies to demonstrate similar stabilities or activities.^14, 22^ Work by Eick et al. has demonstrated that enzyme activity is qualitatively robust to sampling from the probability distribution.^15^ As a result, it is becoming more commonplace to sample ancestral sequences from the probability, either in a biased or unbiased fashion, and demonstrate these resurrected proteins behave similarly to the SMP sequence.^23^ As of yet, a systematic study addressing the hypothesis that the ancestral SMP protein stability is biased has not been explicitly tested.

In a majority of cases the true ancestral state will remain forever hidden from us and so will the true ancestral biochemical property.^24^ Therefore, we cannot directly determine the bias in resurrected SMP sequence property relative to the true ancestral sequence. We are, in fact, only able to experimentally determine if the SMP sequence is biased in activity and stability relative to the average activity and stability of randomly sampled ancestral sequences. The goal of the work presented here is to quantitatively address whether the SMP sequence is biased relative to randomly sampled ancestral sequences. The T_m_ and cognate k_cat_/K_m_ of forty enzymes, four SMP sequences and 36 sampled sequences across four internal nodes, were measured for Apicomplexa lactate and malate dehydrogenases (L/MDH). Overall, we see no bias in T_m_ or k_cat_/K_m_ of the SMP sequence relative to the average activity or stability for randomly sampled sequences. Selecting an SMP sequence for resurrection does not necessarily translate to a stability or activity bias.

## Results

### Apicomplexa L/MDH proteins represent an ideal for statistical analysis

We want to compare the biochemical and biophysical properties of the SMP sequence to many sequences sampled from the probability distribution. *A priori*, we knew we would need to work with a set of enzymes that are easy to express, purify, and characterize. In addition, we wanted a system that would contain evolutionary events that are typically of interest to biochemists and evolutionary biologists alike. For these reasons we chose to work with a system we have studied before, the Apicomplexa L/MDH protein family.^5^

MDHs are essential NAD^+^ dependent oxidoreductases of the TCA cycle and oxidize malate to oxaloacetate. LDH enzymes are structural and functional homologs of MDHs, that reduce pyruvate to lactate under anaerobic conditions regenerating NAD^+^. LDHs have been known to evolve at least four times from ancestral MDHs. Two of these events have occurred in the Apicomplexa phyla and the first instance is along the branch between Anc356 and Anc350, which is shown in a cartoon representing the maximum likelihood phylogenetic tree (fig. 1).

**Fig. 1:**
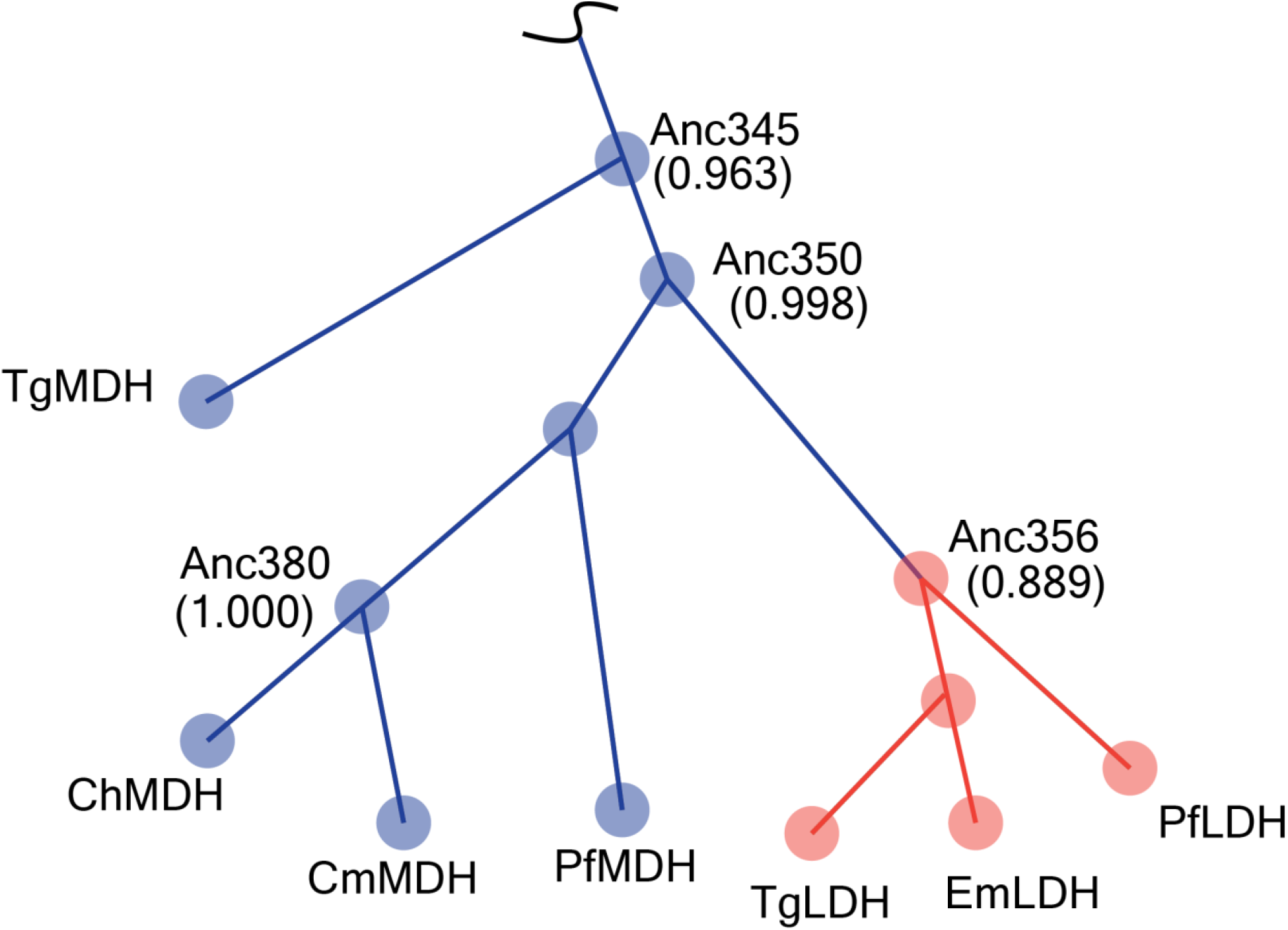
A cartoon representation of the maximum likelihood phylogenetic tree illustrates the evolution of an ancestral LDH enzyme from an ancestral MDH enzyme. The blue indicates malate is the preferred substrate and orange indicate lactate is the preferred substrate. Each terminal and internal node are drawn as filled-in circles represents an extant and ancestral gene respectively. Each node is connected by a branch that is not to scale. The four ancestral sequences resurrected in this study are labelled according to their PAML node number and the corresponding approximate Bayes supports are shown in parentheses near the node PAML node number.

We chose to resurrect Anc356 and Anc350 because they are the result of a transition between to enzymes of different function. Anc356 is the last common ancestor to a group of extant LDHs which evolved from Anc350, an ancestral MDH. We also selected Anc345 because it is the ancestor to all the modern Apicomplexa L/MDH enzymes and was acquired through a horizontal gene transfer event from an ancestral α-proteobacteria. It is possible that the evolutionary events outlined above could have an impact on the precision of our resurrected sequences. Therefore, we chose to resurrect Anc380 because it is a more recent ancestor that is not associated with any change in function or horizontal gene transfer.

### Residue differences between the SMP and sampled sequences are not biased toward any particular domain and are numerous

Previously, our lab has observed that residues outside of the active site of modern Apicomplexa L/MDHs can affect the activity of each enzyme.^5^ We expect that uncertainty throughout the ancestral enzyme sequence will potentially have an impact on enzyme activity and stability. The SMP sequence from each ancestral node was modelled using RoseTTAFold and colored according to the residue probability (supplementary, fig. 1).^10, 25^ In general, the most uncertain residues appear to be surface exposed and not involved in the expected tetramer interface, in the active site, or buried within the protein.

A difference in activity and stability is not necessarily linked to the number of differences between two proteins. However, there is a higher likelihood that two ancestral sequences with only a few amino acid differences are alike as opposed to two ancestral sequences with many amino acid differences. The number of differences between our SMP and sampled sequences range from 15 to 43 (supplementary, Table 8). The average number of differences between the SMP sequence and sampled sequences varied from 21-34. The number of differences between the SMP sequence and sampled sequence were in line with the average probability of the SMP sequence.

### Thermostability of SMP sequence is not biased compared to sampled sequences

Several pieces of literature have noticed that ancestral SMP sequences tend to be more thermostable than their extant counterparts. Therefore, one controversy in the APR field is whether SMP sequences display a thermostability bias as a consequence of the systematically biased method used to generate the sequence.^19–21, 26^ We can determine if the SMP sequence is exceptionally thermostable relative to sequences randomly sampled from the ancestral probability distribution in an unbiased manner. We resurrected 10 sequences per node, 1 SMP sequence and 9 sampled sequences, for four different nodes in our phylogenetic tree and measured their melting temperature by DSC (fig. 2a). The melting temperature for the SMP sequence falls within the range of the melting temperatures measured for the sampled sequences in all instances.

**Fig. 2:**
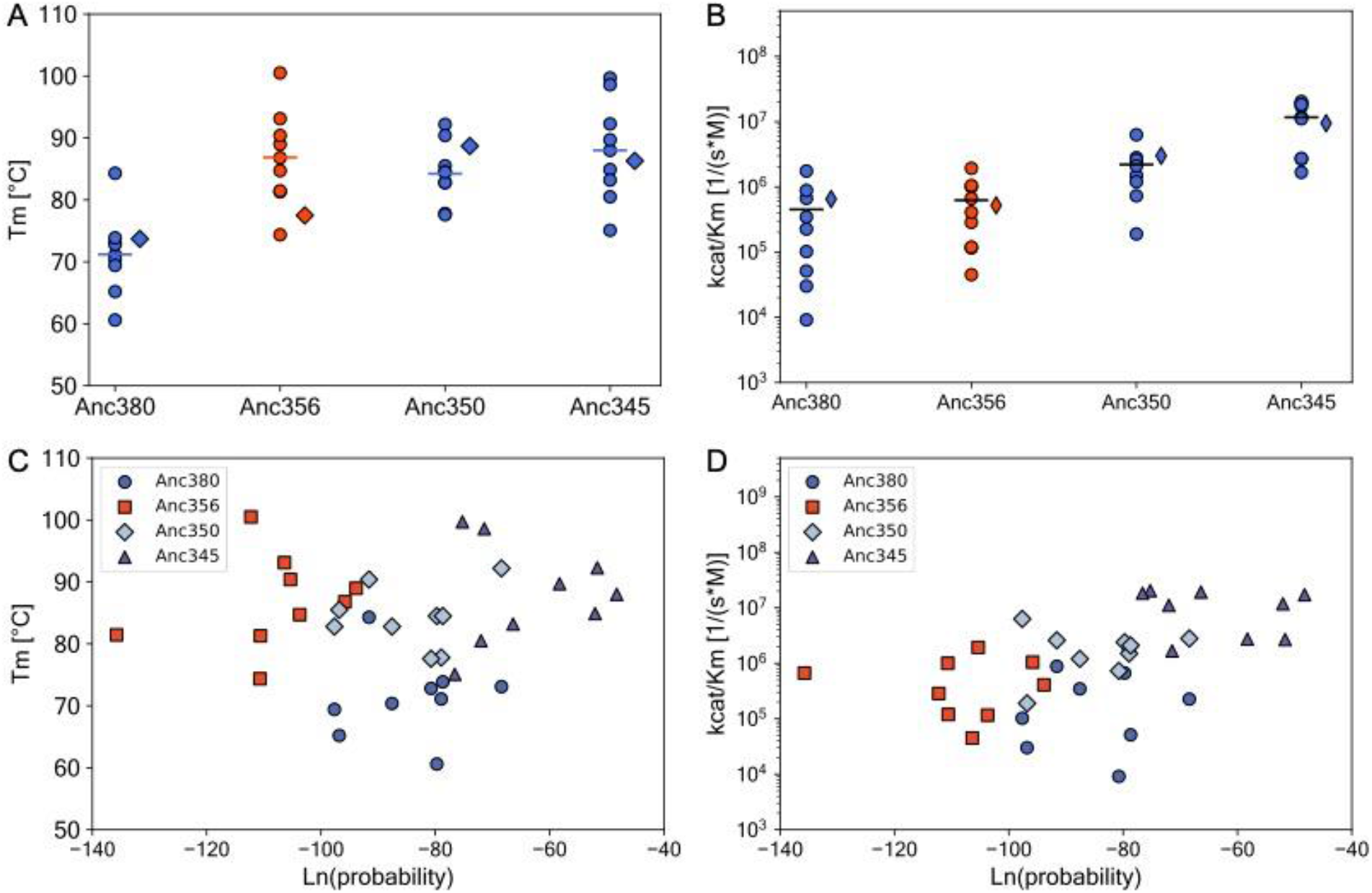
The SMP sequence across all four nodes is not biased in melting temperature or activity compared to the sampled sequences. (*a*) A plot of the melting temperature for each resurrected enzyme for each ancestral node. The melting temperature of the SMP sequence is represented by a ◆, the melting temperature of the alternative sampled sequences by a ●, and the average melting temperature of the alternative sequences by a —. (*b*) The same as (*a*), but for the activity of each protein toward its cognate substrate. (*c*) A scatter plot of each sequence sampled from each ancestral probability distribution as a function of the sequence natural log-probability. (*d*) The same as (*c*), but the activity is plot as a function of natural log-probability.

If the SMP sequence is biased in thermostability then its T_m_ would consistently be above or below the average T_m_ from sequences sampled from the probability distribution. We can test this by subtracting the average sequence T_m_ from the SMP sequence T_m_. If the SMP sequence were systematically biased toward increased thermostability compared to randomly sampled sequences, then we would see ta large positive difference between the average sampled sequence T_m_ and the SMP T_m_. The actual differences are 2.7 °C, −9.5 °C, 4.7 °C, and −1.7 °C for Anc380, Anc356, Anc350, and Anc345 respectively and in fact, the total difference is quite small at −3.8 °C and negative (Supplementary, Table 2).

An extension of the SMP thermostability bias is that some more probable sequences from the ancestral probability distribution are more thermostable than less probable sequences. Incorporating lower probability residues in ancestral sequences could be more detrimental to the protein function overall, because their probabilities are based upon actual substitutions observed in nature.^27, 28^ It is a reasonable hypothesis that the melting temperature of lower probability sequences will tend to be below the average melting temperature, whereas the melting temperature of higher probability sequences will tend to be above the average melting temperature. We test this hypothesis by plotting the protein T_m_ as a function of the sequence log probability (fig. 2c). We observe the melting temperatures are not correlated to sequence log-probability, because the correlation coefficients are 0.02, 0.24, 0.09, and 0.07 for Anc380, Anc356, Anc350, and Anc345 respectively (Supplementary, Table 3).

### Activity of SMP sequence is not biased compared to sampled sequences

In addition to a thermostability bias, a bias in enzyme activity has been suggested for SMP sequences. To test this hypothesis, we measured the activity (*k*_cat_/K_m_) of each resurrected SMP sequence and ancestors sampled from the probability distribution. We plot the activity on a log_10_-scale for each enzyme across all four ancestral nodes, as well as the average activity of the sampled ancestral sequences (fig. 2b). We report log-activity as a matter of convenience because it allows us to see a wide value range on one figure. The difference in log-activity between each SMP sequence and average of the alternative sequences for all nodes is only −0.15 (Supplementary, Table 4).

We also sought to determine if there was any preference for a low probability sequence to display low activity. We plot enzyme log-activity as a function of sequence log-probability and found no correlation (fig. 2d). The correlation coefficients are 0.17, 0.05, −0.21, and −0.41 for Anc380, Anc356, Anc350, and Anc345 respectively (Supplementary, Table 5).

The activity is determined by how fast the enzyme can conduct its chemical reaction at saturating substrate concentration (k_cat_) and its substrate concentration at one half k_cat_ (K_m_). It is possible that possible that the individual parameters of the activity for each SMP sequence are actually biased, but these biases cancel for the activity. We also plot the k_cat_ and K_m_ of each resurrected enzyme (supplementary, fig. 2). Except for the individual k_cat_ for Anc380, all average sequence k_cat_ values and SMP sequence k_cat_ values are close to 0. For Anc380, the SMP sequence is nearly 80 s^−1^ slower than the average k_cat_. However, the within range variability is randomly sampled sequence k_cat_ s is greater than the difference between the average and the SMP k_cat_ (Supplementary, Table 6).

Similarly, the individual K_m_ values of each SMP sequence falls within the range of K_m_ values for sampled sequences. The total difference between natural log-*K_m_* of the SMP and average sampled sequences is only −0.08 (supplementary, Table 7). The SMP sequence has a slightly lower K_m_ than the average sequence.

### Isoelectric point and extinction coefficient of SMP sequence may be biased compared to sampled sequences

No known function exists to accurately estimate the thermostability or activity of an enzyme, so their values had to be measured. Consequently, we were limited by the number of proteins we could physically resurrect. A sample size of 9 may not be sufficient to observe a bias. If we could sample and measure the T_m_ or activity of 100, 1,000, or 10,000 proteins we may find that many proteins low probability proteins display low melting temperatures or no activity. We would like to increase our sample size to determine if our results are robust sampling error.

To increase our sample size, we turn to properties we can calculate relatively accurately using computational approaches. For example, we can accurately estimate the isolectric point (pI) and extinction coefficient (ε) of enzymes, and we can also calculate the molecular weight exactly. We calculated the pI, ε, and molecular weight of 10,000 sequences sampled from each ancestral probability distribution. We see that the pI, ε, and molecular weight for the SMP sequence lie within one standard deviation of the mean, except for the pI of Anc356 which lies within two standard deviations (fig. 3 and Supplementary, Table 9). Importantly, the SMP pI and ε values are consistently above and below their respective average pI and ε values (fig. 3a and 3b). This consistency points toward a subtle bias in the SMP sequence as compared to the average sequence.

**Fig. 3:**
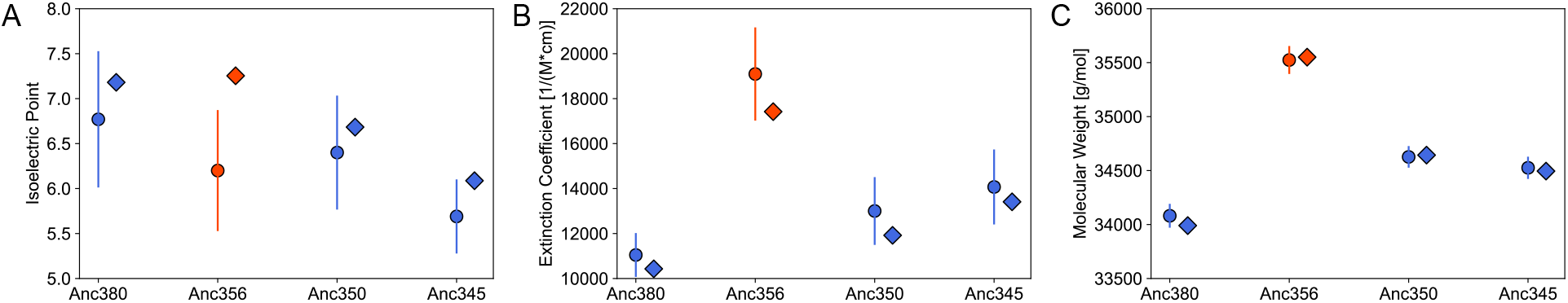
Theoretical and calculated properties of SMP sequence may be biased compared to the average property of sampled sequences. A plot of the pI, extinction coefficient, and molecular weight enzymes sampled from the probability distribution. The circles represent the average of 10,000 sampled sequences and the error bars are the standard deviation. The diamonds represent the SMP sequence.

### No trade-off between stability and activity

Recently, ASR has gained attention as a potential protein engineering tool, because of the purported exceptional thermostability of ancestral SMP sequences. There is a hypothesis that increasing the protein stability will correspondingly decrease the enzyme activity. We can test this hypothesis with our ancestral dataset by plotting the melting temperature against the activity of each protein (fig. 4). The median melting temperature among all enzymes is 84 °C and the median activity is 10^6^ s^−1^*M^−1^. We find that, among our resurrected proteins, there are enzymes with both melting temperatures and activities above the median values. The correlation coefficient for the data is 0.24.

**Fig. 4:**
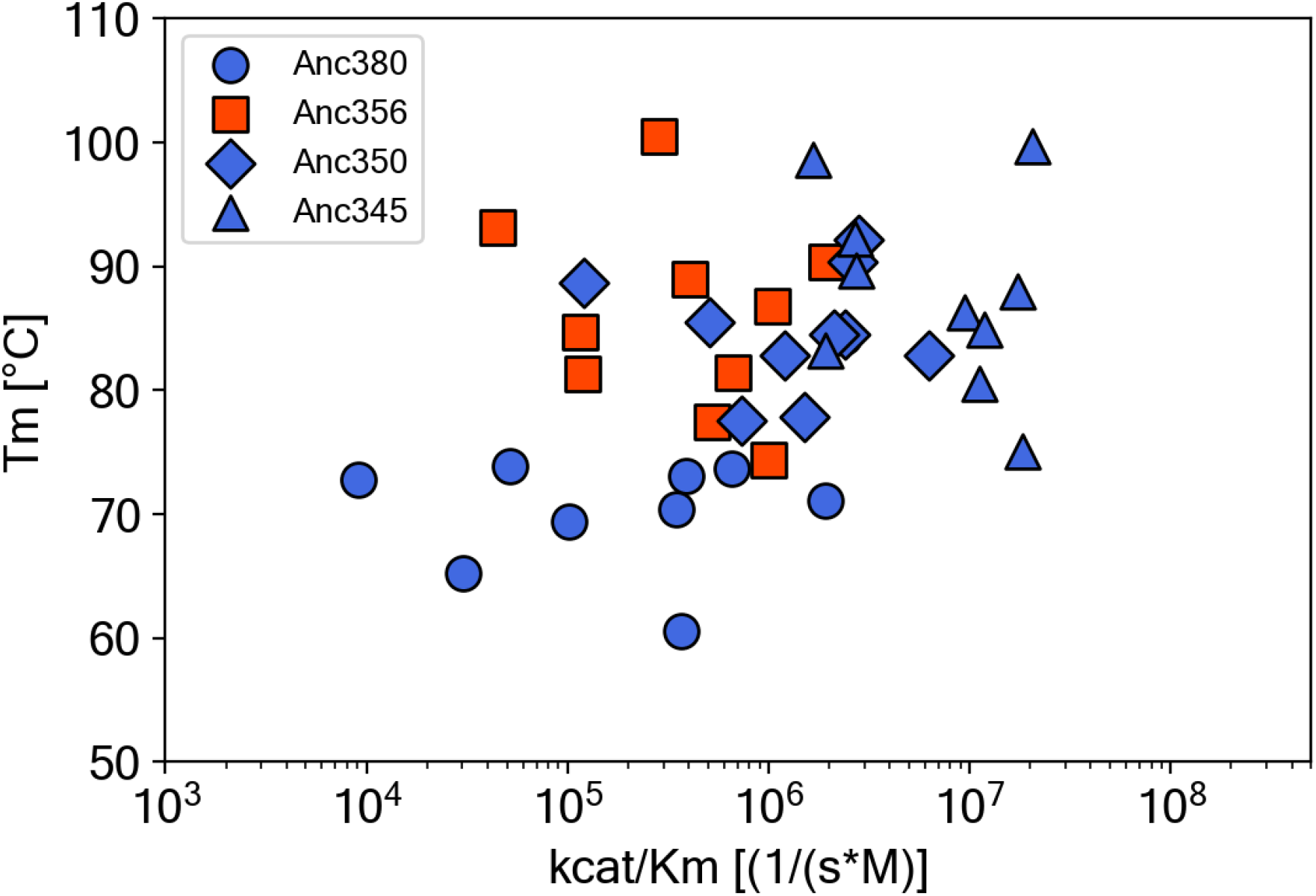
The melting temperature and activity of resurrected enzymes are not correlated. A plot of the T_m_ as a function of log-activity for each resurrected enzyme. Ancestral LDH enzymes are in orange-red and MDH enzymes are in blue.

## Discussion

APR is a biochemical tool used elucidate the origins of extant protein function and structure by resurrecting ancestral proteins to study *in vitro*. Ancestral proteins are reconstructed by selecting the most likely residue from a probability distribution of amino acids at each site of the protein. This is a systematically biased method of generating an ancestral sequence. Does selecting the most probable residue at each site of an ancestral sequence result in a protein having a physicochemical bias? We have answered this question by measuring the properties of ancestral sequences generated in an unbiased way and comparing their average property value to the corresponding SMP sequence property value. We see that the physicochemical properties of the SMP sequences is not biased in comparison to randomly sampled sequences of the probability distribution.

### The SMP sequence T_m_ is an unbiased representative of the posterior probability distribution

An amino acid probability distribution is calculated at each site of an ancestral sequence in APR. This means that there are an astronomical number of plausible ancestral sequences that could be resurrected. To circumvent this problem an ancestral sequence is generated by selecting the most probable residue at each site. Recently, we have shown that the SMP sequence is actually the sequence with the most expected correct residues and thus could be investigated as the single best estimate of the true ancestral sequence. However, this does not preclude the SMP sequence from exhibiting a systematic bias in biochemical properties because selecting the SMP residue at each site is a systematically biased method to generate an ancestral sequence.

Indeed, in a computational analysis Williams et al. calculated that SMP ancestral sequences have a stability bias in comparison to the true sequence. This bias was reduced when sequences were randomly sampled from the ancestral probability distribution. Unfortunately, with real data we do not know the true ancestral sequence. To experimentally assess if the T_m_ of the SMP ancestral sequence is a biased estimate of the ancestral probability distribution we need to measure an unbiased estimate.

An unbiased method of choosing an ancestral sequence is to randomly sample them from the posterior probability distribution. Therefore, sampling several of these sequences, measuring their biochemical property, and taking the average of that property should give an unbiased estimate of the biochemical properties of the probability distribution. This is essentially an experimental Monte Carlo sampling method to determine the biochemical property of a distribution. We performed this Monte Carlo sampling method by measuring the T_m_ of sequences sampled from the posterior probability distribution of four different ancestral nodes in an L/MDH phylogeny. The difference between the average sequence and SMP sequence T_m_ is near 0 for all ancestors, with some differences being greater than 0 and others less than 0. Therefore, the SMP ancestral sequence T_m_ is an unbiased point estimate of the average ancestral sequence T_m_.

In addition to a stability bias, Williams et al. suggest that the bias would extend to other properties like enzyme activity. We also measured the activity of each ancestral sequence, including the individual k_cat_ and K_m_, and found that that the difference between the average ancestral sequence activity, k_cat_, and log-K_m_ and the corresponding SMP ancestral sequence values were near 0 for most measurements. The only exception is the k_cat_ for Anc380. However, the standard deviation in the k_cat_ of Anc380 is larger than the difference between the SMP and average value. The SMP ancestral sequence is generally representative of activity of an ancestral distribution.

### The pI and ε of the SMP sequence are systematically biased estimates of the posterior probability distribution

An experimental Monte Carlo sampling approach to approximate the expected property of a distribution limited the number of samples we could experimentally characterize. Increasing the sample size may reveal a systematic bias in the T_m_ or activity of SMP ancestral sequences. To address this sample size problem we calculated the average values of other biophysical properties of proteins like isoelectric point, extinction coefficient, and molecular weight from 10,000 samples and compared them to the values of the SMP ancestral sequence. Like with T_m_ and activity, we still find that the individual differences between the average ancestral sequence and SMP ancestral sequence pI, ε, and molecular weight are less than 1 standard deviation for all calculations except the pI of Anc356. However, unlike with T_m_ and activity we find that the SMP ancestral sequence pI is consistently above the average sequence pI and the SMP ancestral sequence ε is consistently below the average sequence ε. While the SMP ancestral pI and ε are good estimates, they appear to be slightly biased.

### There is no evidence for systematic bias in more probable ancestral sequence melting temperatures

Even though the properties of the SMP sequence is representative of the average sequence sampled from the probability distribution, it does not mean there is no systematic bias from usually including the most probable residue. It is reasonable to hypothesize that the stability of a protein is a function, to some degree, of the probability of a sequence. For example, if we generated a sequence by selecting the single least probable residue from the ancestral probability distribution then we will expect that protein to be exceptionally unstable if it folds at all. If this hypothesis holds, then the more probable a sequence is the more stable it is likely to be.

We tested this hypothesis by plotting the melting temperature of all 36 sampled proteins as a function of sequence log-probability. The melting temperature is not a function of the sequence log-probability, because the correlation coefficients are all less than an absolute value of 0.24. For our sample size, a more probable sequence does not necessarily result in a protein with a greater thermostability. The analysis was repeated with the protein activity and a similar result was found. Therefore, there is no evidence for systematic bias in thermostability and activity when selecting more likely enzymes to resurrect. The results may change as more samples are acquired because the extremes of the probability distribution will be sampled.

### The SMP sequence is not the most stable enzyme in the probability distribution

Previous studies have attempted to address the origin of the purported SMP sequence stability bias. Williams et al. took a computational approach and simulated sequences along a phylogeny, calculated the stability of the true ancestral sequence, SMP ancestral sequence, and sampled ancestral sequence. The sampled ancestral sequences were closer in stability to the true sequence than the SMP ancestral sequence. The hypothesis is that sampled sequences are less stable because there are more detrimental incorrect mutations as a result of incorporating lower probability residues. The authors suggest that enzyme activity could mirror their observations on stability.

Similarly, an experimental approach by Trudeau et al. argues that SMP ancestral sequences are exceptionally stable because of the “consensus effect”. This argument is based on the empirical observation that mutating an uncommon amino acid to the most common amino acid tends to stabilize a protein. The SMP sequence generally contains some of the most common amino acids at different positions, potentially resulting in the most stable protein. This is essentially the same argument as Williams et al., but from the opposite perspective.

The implication from those two studies is that the SMP sequence should, more often than not, be the most stable resurrected protein. However, from our T_m_ measurements we find that incorporating less likely (*i.e*. more uncommon) amino acids in a reconstruction can actually result in significantly more stable proteins compared to always incorporating the SMP residue. Similarly, we find that incorporating less likely amino acids can increase ancestral enzymes activity. This could be a result of epistasis in a protein sequence, a potentially uncommon amino acid in one sequence context that reduces stability could in fact increase stability in another sequence context.

### There is substantial variability in the T_m_ of sampled ancestral resurrections

APR has been used to infer properties of the environment in which ancestral enzymes existed in. However, any one sequence is highly improbable and there are many plausible ancestral sequences. On top of that biochemists know that a single mutation can have a large impact on the stability of a protein. For this reason, experimentalists will resurrect alternative ancestral sequences from the ancestral probability distribution to determine if their results are robust to sequence uncertainty. In one example, Gaucher et al. resurrected 5 additional sequences randomly sampled from an ancestral probability distribution to demonstrate that one SMP sequence was not a biased estimate of the temperature for precambrian earth.^3^ The standard deviation of the T_m_ for the last common ancestor studied was ~2.7 °C.

Often, however, a sequence is not randomly sampled from the probability distribution. Rather, a sequence or set of sequences from the ancestral probability distribution are restricted to residues above an arbitrary probability threshold. Hart et al. resurrected 10 ancestral sequences, in addition to the SMP sequence, excluding probabilities below 0.2, to demonstrate the trends they observed were not simply a function of sequence uncertainty.^23^ The standard deviation of their last common ancestor studied was ~1.6 °C. This deviation is likely to be a function of the type of protein studied and the evolutionary model used.

We have resurrected 9 samples across four nodes of varying depth in a phylogenetic tree and find that the standard deviation of the T_m_ for the sampled sequences exhibited one standard deviation that ranged between 5-8 °C. This is elevated compared to previous studies, but we have sampled more sequences and did not constrain residue probability. These standard deviations we observed reduce our confidence in any trend or bias based on ancestral sequence thermostability. Given the spread of the T_m_ measurements it is difficult to determine if ancestors associated with our phylogeny are actually more thermostable than today’s extant sequences.

### There is no stability-activity trade-off in our ancestral resurrections

The stability of an enzyme and its activity are two characteristics that are not necessarily mutually independent of one another. A residue needed for activity may destabilize the enzyme and a residue which may stabilize the enzyme could reduce activity. Several studies have investigated and found a stability-activity trade-off between substitutions made in different enzymes.^29, 30^ However, these studies investigate the trade-off between a handful of residues. We would like to know if this trade-off is observed for many substitutions of residues outside of the active site of an enzyme.

Our L/MDH ancestral resurrections sampled from the ancestral probability distributions vary from the SMP sequences at 15-43 sites. Each sequence also varies in activity and stability and we found that ancestral sequences can exhibit exceptional stability and activity. There exist sequences that are both highly active and stable. For example, a sampled sequence from the ancestral probability distribution of Anc345 is the most active enzyme and has a T_m_ near 100 °C. Sampling ancestral enzyme sequences is a viable protein engineering route to generate enzymes with industrially relevant properties, like high stability or activity.

Analyses have also pointed toward a trade-off between developing a new function and the stability of a protein.^31^ In our Apicomplexa L/MDH phylogeny we have an example of one enzyme evolving a new function. Two mutations in the active-site of the ancestral LDH (Anc356), a lysine to arginine mutation and a five amino acid insertion, are responsible for the change in function from an ancestral MDH (Anc350). In addition to the thermal stability independence of activity within a node, we observe that the stability for Anc356 is not impaired after evolving a change in substrate specificity from malate to lactate by active-site modification (fig. 2). In fact, we see the opposite, the average melting temperature of Anc350 is 84.2 °C and the average melting temperature of Anc356 is 86.8 °C.

## Conclusion

We have demonstrated experimentally that although the SMP sequence is generated in a biased fashion there is no observable bias in stability or function of the enzyme relative to randomly sampled alternative ancestors. The SMP sequence T_m_ and activity is representative of the probability distribution. Therefore, we suggest that, when using ASR as a tool, the SMP sequence is a sufficient representative sequence for inferring ancestral properties. In addition, we do not see any correlation between activity and stability. If a resurrected sequence is highly active there is no reason to assume that the stability will be diminished. Inferences drawn from one property may not provide information on any other property. This is useful for protein engineering purposes because randomly sampling ancestral sequences may result in highly stable and highly active enzymes.

## Materials & Methods

### Ancestral sequence reconstruction

Previously our lab reported on the construction of a maximum likelihood phylogenetic tree used to resurrect ancestral sequences.^5, 10, 11^ The alternative ancestral sequences were sampled according to the weighted probability of each amino acid at each site. For each node of interest 9 randomly sampled sequences were selected for testing.

### Protein Expression & Purification

A pET-24 a (+) vector containing each codon optimized His-tagged sequence was ordered (Genscript) and transformed into *E. coli* BL21 (DE3) pLysS cells (Novagen). Cells were grown to an O.D._600_ of 0.6 at 37 °C, induced with 500 μM IPTG, and expressed for four hours at 37 °C. Cells were pelleted and stored at −20 °C for future use. Pelleted cells were resuspended and lysed by sonication. The soluble lysate was separated from insoluble lysate by centrifugation at 25,000 x g for 30 minutes. The supernatant was decanted and filtered to 0.22 μ and purified using an imidazole gradient on a HisTrap™ HP column (GE Healthcare Life Sciences). The purified protein was concentrated in Amicon® Ultra-15 centrifugal filter units (MilliporeSigma) buffer exchanged into storage buffer (50 mM Tris-Base, 100 mM NaCl, 0.1% EDTA, 0.02% sodium azide, 300 μM TCEP-HCl, pH 7.4) using a PD-10 desalting column (GE Healthcare Life Sciences).

### Steady-state methods

Enzyme and NADH cofactor are held constant as substrate (oxaloacetic acid or sodium pyruvate) is varied. Each enzyme is diluted from storage buffer to assay buffer (50 mM Tris-Base, 50 mM NaCl, pH7.4) in order to measure activity. Final enzyme concentrations ranged from 1 nM to 1 μM varied depending on individual enzyme turnover rates. The NADH concentration was maintained at 200 μM across all enzyme constructs tested. The decrease in NADH absorbance at 340 nm was monitored over time on the Cary 100 Bio or SX-20 Stopped-Flow at 25 °C. The kinetic parameters for each enzyme were estimated from a chi-squared to the Michaelis-Menten equation 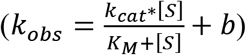 or the partial substrate model of inhibition 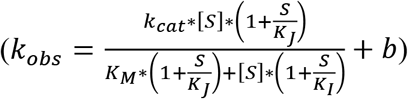 in KaleidaGraph.

### Melting temperature methods

The melting temperature was determined by increasing the temperature from 25-110 °C and monitoring the difference in heat required to raise the sample cell with protein compared to the reference cell with buffer on a Nano DSC (TA Instruments, New Castle, DE). The protein sample was buffer exchanged into 50 mM K_2_HPO_4_, pH 7.5, 50 mM KCl by SEC or dialysis.

### Theoretical pI, ε, and molecular weight calculation

Following the protocol and C+ script of Kozlowski the pH at which the sequence has no charge was calculated in Python.^32^ Briefly, an empirically derived pKa for each charged residue, as well as the N- and C-terminus of the sequence are tallied. Using a bisection algorithm, the pH at which the protein has no charge (*i*.*e*. the pI) is determined to a resolution of 0.01 with the Henderson-Hasselbach equation under reducing conditions. The pI was then calculated for the SMP sequence and 10,000 sequences sampled from the probability distribution. The average and standard deviation were calculated for the pI of the 10,000 samples. This process was repeated for the extinction coefficient and molecular weight, which were calculated using the ProtParam module of Biopython. ^33^

### Ancestral sequence structure and visualization

The structure of each SMP sequence was predicted using RoseTTAFold and visualized using PyMol.^25, 34^ The posterior probability of each SMP residue was extracted from the CODEML rst file and mapped onto the structure and colored according to probability.

## Notes

### Competing Interest Statement

The authors have declared no competing interest.

