## Supplemental for "The most probable ancestral sequence reconstruction yields proteins without systematic bias in thermal stability or activity"

### Supplemental Figures

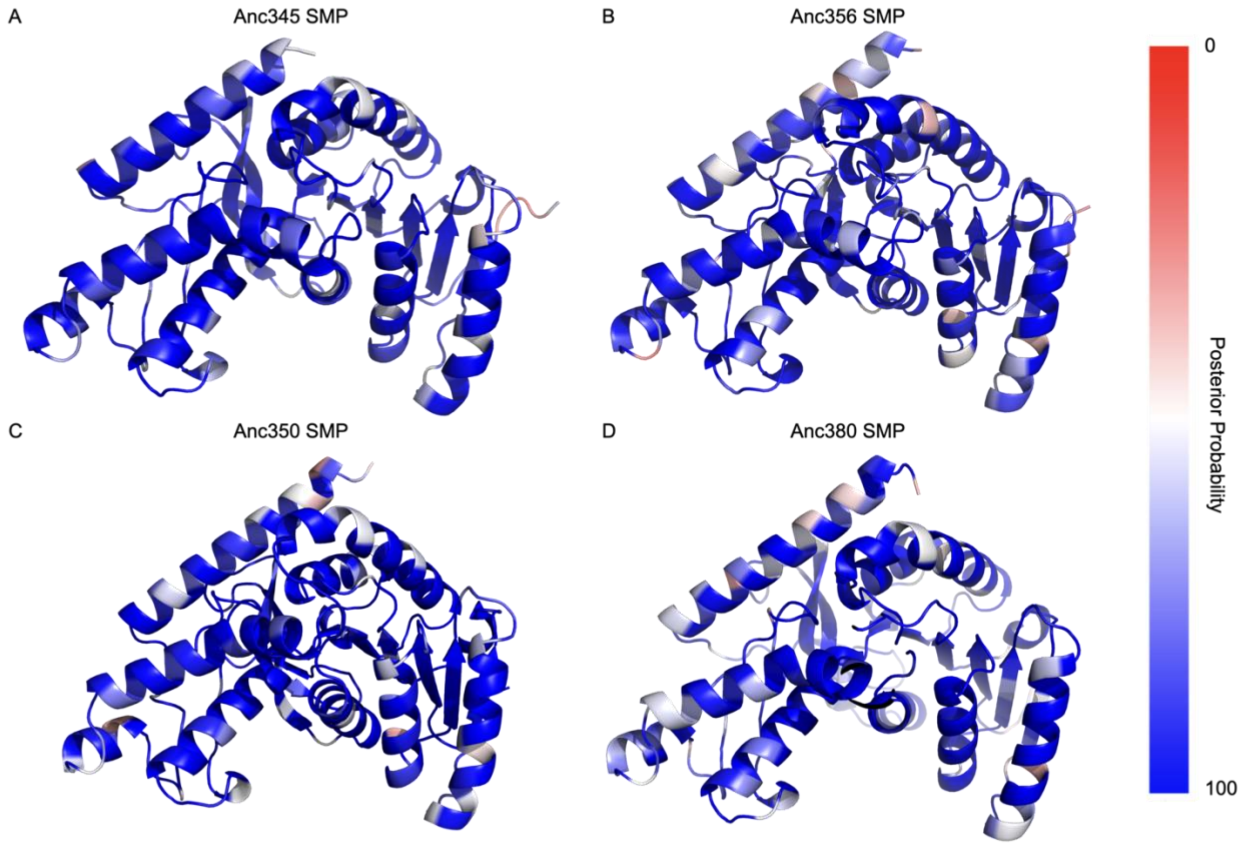


SI Fig. 1: **Uncertain sites are distributed throughout the surface of each enzyme.** The residue in each sequence is colored by posterior probability. Blue is the most certain whereas red is the most uncertain residue.

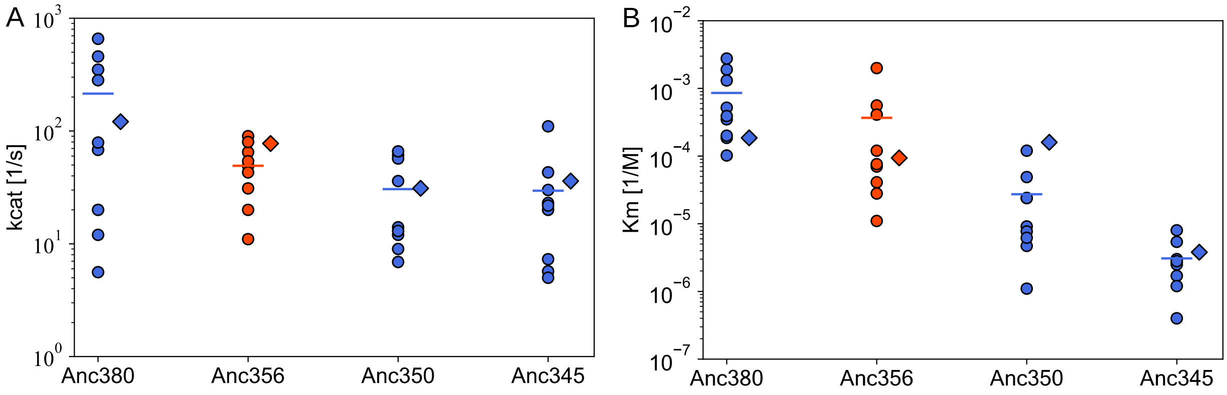


SI Fig. 2: **The SMP sequence $k_{cat}$ and $K_m$ are close to the average values of sampled sequences.** (*a*) A plot of the $k_{cat}$ for each enzyme resurrected across all four nodes. The $k_{cat}$ of the SMP sequence is represented by a ◆, the $k_{cat}$ of the alternative sampled sequences by a ○, and the average $k_{cat}$ of the alternative sequences by a —. (*b*) is the same as (*a*), but with the $K_m$ rather than the $k_{cat}$.

| Sequence | Anc380 | Anc356 | Anc350 | Anc345 |
|---|---|---|---|---|
| SMP | -39.260 | -53.908 | -40.157 | -31.302 |
| S1 | -96.806 | -106.390 | -82.679 | -52.050 |
| S2 | -79.732 | -103.720 | -83.122 | -66.392 |
| S3 | -78.961 | -110.670 | -90.826 | -75.257 |
| S4 | -68.421 | -112.250 | -77.494 | -72.002 |
| S5 | -91.599 | -93.856 | -83.058 | -48.262 |
| S6 | -87.574 | -110.590 | -65.760 | -71.433 |
| S7 | -97.645 | -105.350 | -89.373 | -51.665 |
| S8 | -80.740 | -95.827 | -73.525 | -58.264 |
| S9 | -78.687 | -135.750 | -86.713 | -76.598 |

SI Table 1: A table summarizing the log-probability of each enzyme resurrected in this study from figure 2.

| Sequence | Anc380 | Anc356 | Anc350 | Anc345 |
|---|---|---|---|---|
| SMP | 73.7 | 77.5 | 88.7 | 86.3 |
| S1 | 65.2 | 93.1 | 85.5 | 84.9 |
| S2 | 60.6 | 84.7 | 84.5 | 83.2 |
| S3 | 71.1 | 74.4 | 77.8 | 99.7 |
| S4 | 73.1 | 100.5 | 92.2 | 80.5 |
| S5 | 84.3 | 88.9 | 90.4 | 88.0 |
| S6 | 70.4 | 81.3 | 82.8 | 98.6 |
| S7 | 69.4 | 90.4 | 82.8 | 92.3 |
| S8 | 72.8 | 86.8 | 77.6 | 89.7 |
| S9 | 73.9 | 81.5 | 84.5 | 75.1 |
| Avg | 71 (6) | 87 (8) | 84 (5) | 88 (8) |
| SMP-Avg | 2.7 | -9.5 | 4.7 | -1.7 |

SI Table2: Summary of melting temperature data in Celsius from figure 2.

| | slope | intercept | R |
|---|---|---|---|
| Anc380 | 0.01 | 72.1 | 0.02 |
| Anc356 | 0.15 | 103.3 | 0.24 |
| Anc350 | 0.05 | 88.1 | 0.09 |
| Anc345 | 0.05 | 91.2 | 0.07 |

SI Table 3: Summary of line of best fit for $T_m$ vs. log-probability plots from figure 2.

| Sequence | Anc380 | Anc356 | Anc350 | Anc345 |
|---|---|---|---|---|
| SMP | 5.8 | 5.7 | 5.3 | 7.0 |
| S1 | 4.5 | 4.7 | 5.7 | 7.1 |
| S2 | 5.8 | 5.1 | 6.4 | 7.3 |
| S3 | 6.2 | 6.0 | 6.2 | 7.3 |
| S4 | 5.4 | 5.5 | 6.4 | 7.0 |
| S5 | 5.9 | 5.6 | 6.4 | 7.2 |
| S6 | 5.5 | 5.1 | 6.1 | 6.2 |
| S7 | 5.0 | 6.3 | 6.8 | 6.4 |
| S8 | 4.0 | 6.0 | 5.9 | 6.4 |
| S9 | 4.7 | 5.8 | 6.3 | 7.3 |
| Avg | 5.7 | 5.8 | 5.8 | 7.1 |
| SMP-Avg | 0.1 | -0.1 | -0.5 | -0.1 |

SI Table 4: A table summarizing the log-activity from figure 2.

| | slope | intercept | R |
|---|---|---|---|
| Anc380 | 1.03E4 | 1.32E6 | 0.17 |
| Anc356 | 2.57E3 | 9.02E4 | 0.05 |
| Anc350 | -3.81E4 | -1.01E6 | -0.21 |
| Anc345 | -1.98E-5 | -8.86E5 | -0.29 |

SI Table5: Summary of line of best fit for Activity vs. log-probability plots from figure 2.

| Sequence | Anc380 | Anc356 | Anc350 | Anc345 |
|---|---|---|---|---|
| SMP | 121 | 49 | 31 | 36 |
| S1 | 5.6 | 90 | 61 | 20 |
| S2 | 68 | 65 | 57 | 23 |
| S3 | 350 | 11 | 14 | 110 |
| S4 | 79 | 20 | 66 | 30 |
| S5 | 458 | 31 | 12 | 43 |
| S6 | 658 | 49 | 9 | 5 |
| S7 | 282 | 54 | 6.9 | 7.5 |
| S8 | 12 | 43 | 36 | 22 |
| S9 | 20 | 80 | 13 | 7 |
| Avg | 200 (200) | 50 (30) | 30 (20) | 30 (30) |
| SMP-Avg | -79 | 1 | 1 | 6 |

SI Table 6: A table summarizing $k_{cat}$ in units of $s^{-1}$ from SI figure 3.

| Sequence | Anc380 | Anc356 | Anc350 | Anc345 |
|---|---|---|---|---|
| SMP | -3.7 | -4.0 | -3.8 | -5.4 |
| S1 | -3.7 | -2.7 | -3.9 | -5.8 |
| S2 | -4.0 | -3.3 | -4.6 | -5.9 |
| S3 | -3.7 | -5.0 | -5.0 | -5.3 |
| S4 | -3.5 | -4.2 | -4.6 | -5.6 |
| S5 | -3.3 | -4.1 | -5.3 | -5.6 |
| S6 | -2.7 | -3.4 | -5.1 | -5.5 |
| S7 | -2.6 | -4.6 | -6.0 | -5.6 |
| S8 | -2.9 | -4.4 | -4.3 | -5.1 |
| S9 | -3.4 | -3.9 | -5.2 | -6.4 |
| Avg | -3 (3) | -3 (3) | -5 (4) | -6 (6) |
| SMP-Avg | -0.7 | -0.7 | 1.2 | 0.6 |

SI Table 7: A table summarizing log-$K_m$ with $K_M$ in units of M from SI figure 3.

| Sequence | Anc380 | Anc356 | Anc350 | Anc345 |
|---|---|---|---|---|
| SMP | 0 | 0 | 0 | 0 |
| S1 | 35 | 34 | 34 | 16 |
| S2 | 24 | 30 | 32 | 20 |
| S3 | 33 | 35 | 29 | 29 |
| S4 | 20 | 38 | 33 | 27 |
| S5 | 33 | 29 | 25 | 15 |
| S6 | 26 | 34 | 25 | 22 |
| S7 | 33 | 31 | 27 | 18 |
| S8 | 27 | 32 | 23 | 19 |
| S9 | 26 | 43 | 35 | 26 |
| Avg | 29 | 34 | 29 | 21 |

SI Table 8: A table the number of differences between the SMP reconstruction and the sequences sampled from the ancestral probability distribution.

| | pI | ε ($M^{-1}cm^{-1}$) | Molecular Weight (g/mol) |
|---|---|---|---|
| Anc380 SMP | 7.2 | 10430 | 33990 |
| Anc380 Avg | 6.8 (0.8) | 11000 (1000) | 34100 (100) |
| Anc380 SMP-Avg | 0.4 | -570 | -110 |
| Anc356 SMP | 7.3 | 17420 | 35551 |
| Anc356 Avg | 6.2 (0.7) | 19000 (2000) | 35500 (100) |
| Anc356 SMP-Avg | 1.1 | -1580 | 51 |
| Anc350 SMP | 6.7 | 11920 | 34643 |
| Anc350 Avg | 6.4 (0.6) | 13000 (2000) | 34600 (100) |
| Anc350 SMP-Avg | 0.3 | -1080 | 43 |
| Anc345 SMP | 6.1 | 13410 | 34495 |
| Anc345 Avg | 5.7 (0.4) | 15000 (2000) | 34500 (100) |
| Anc345 SMP-Avg | 0.4 | -1590 | -5 |
| Total Difference | 2.2 | -4820 | -21 |

SI Table 9: A table summarizing the theoretical calculations using Python from figure 3.
